# White matter hyperintensities and neuropsychiatric symptoms in Alzheimer’s disease and mild cognitive impairment

**DOI:** 10.1101/791657

**Authors:** Karen Misquitta, Mahsa Dadar, D. Louis Collins, Maria Carmela Tartaglia, Alzheimer’s Disease Neuroimaging Initiative

## Abstract

Background and Purpose: Neuropsychiatric symptoms (NPS) are frequently encountered in patients with Alzheimer’s disease (AD). Focal grey matter atrophy has been linked to NPS development. Cerebrovascular disease can cause focal lesions and is common among AD patients. As cerebrovascular disease can be detected on MRI as white matter hyperintensities (WMH), this study evaluated WMH burden in mild cognitive impairment (MCI), AD and normal controls and determined their relationship with NPS. Methods: NPS were assessed using the Neuropsychiatric Inventory and grouped into subsyndromes. WMH were measured using an automatic segmentation technique and mean deformation-based morphometry was used to measure atrophy of grey matter regions. Results: WMHs and grey matter atrophy both contributed significantly to NPS subsyndromes in MCI and AD subjects, however, WMH burden played a greater role. Conclusions: This study could provide a better understanding of the pathophysiology of NPS in AD.

## 1. Introduction

A majority of individuals diagnosed with Alzheimer’s Disease (AD) also suffer from neuropsychiatric symptoms (NPS). NPS can reduce quality of life, contribute to caregiver burden and lead to institutional care(Kaufer et al., 1998). Moreover, NPS can be difficult to treat(Ryu, Katona, Rive, & Livingston, 2005). Some studies suggest that NPS may worsen cognitive symptoms and functional decline and have associated these symptoms with accelerated mortality(Palmer et al., 2010). Recent work has attempted to identify the incidence and prevalence of these symptoms during the progression of AD(Constantine G. Lyketsosa, Maria C. Carrillob, J. Michael Ryanc, Ara S. Khachaturiand, Paula Trzepacze, Joan Amatniekf, Jesse Cedarbaumg, Robert Brashearh & Milleri, 2012). However, very little is known about the underlying pathophysiology and neuroanatomical correlates of NPS in AD.

NPS are known to exist even in those with mild cognitive impairment (MCI)(Constantine G. Lyketsosa, Maria C. Carrillob, J. Michael Ryanc, Ara S. Khachaturiand, Paula Trzepacze, Joan Amatniekf, Jesse Cedarbaumg, Robert Brashearh & Milleri, 2012; Geda et al., 2008). They have been found to occur before cognitive decline(Lanctôt et al., 2017), and some studies have suggested that specific NPS may be useful as early predictors of AD or dementia(Geda et al., 2008). NPS have also predicted faster progression from MCI to AD(Peters et al., 2015). Moreover, cognitive impairment and NPS may have distinct neuroanatomical deficits in AD(Bruen, McGeown, Shanks, & Venneri, 2008; Shinno et al., 2007). For example, abnormalities in the anterior cingulate cortex were observed in AD patients with delusional thinking compared to those without delusions, and these markers were found to be unrelated to cognition(Shinno et al., 2007).

Reduction in grey matter (GM) volume in discrete cortical and subcortical regions has been associated with specific NPS in AD patients. Bruen et al.(Bruen et al., 2008) used voxel-based morphometry (VBM) to evaluate differences in regional grey matter density associated with NPS in mild AD. They found delusions, agitation and apathy related to cortical atrophy, particularly in the right hemisphere compared to the left, and in the anterior region. Irritability, anxiety and aberrant motor behaviour have been related to atrophy of the amygdala in early AD(Poulin, Dautoff, Morris, Barrett, & Dickerson, 2011), while apathy has been related to atrophy in the dorsolateral and medial prefrontal cortex, anterior cingulate areas, and the caudate and putamen(Bruen et al., 2008; Hahn et al., 2013). These studies suggest that NPS, particularly depression, apathy and delusions, are most frequently associated with changes in the frontal and subcortical regions of the brain. Nevertheless, atrophy alone has not been sufficient to account for NPS in AD and other dementias(Berlow et al., 2010).

White matter hyperintensities (WMH) are white matter lesions in the brain that appear as high signal intensity regions on T2-weighted MRI. They have a number of possible pathological substrates including blood-brain barrier leakage, hypoperfusion, ischemia/hypoxia, inflammation, neurodegeneration and amyloid angiopathy(Gouw et al., 2011). WMHs that result from small vessel disease (SVD) have been associated with vascular risk factors, like hypertension(Shim et al., 2015). WMH load has been associated with AD pathology(Alosco et al., 2018; Dadar et al., 2018), and a relationship between SVD and WMH lesions in AD has been reported in other studies(Shim et al., 2015). WMHs of presumed vascular origin have also been associated with NPS in various populations, including AD(Berlow et al., 2010; Dadar, Maranzano, et al., 2017). However, there is limited research on the contribution of WMH burden to changes in NPS over time in patients with MCI and AD.

The purpose of this study was to evaluate WMH burden and regional GM atrophy in MCI and AD and determine their contribution to NPS over time, using longitudinal data from a large multi-center database from the Alzheimer’s Disease Neuroimaging Initiative (ADNI).

## 2. Materials and Methods

Anonymized data and materials have been made publicly available at the ADNI repository and can be accessed at http://adni.loni.usc.edu/data-samples/access-data/.

### 2.1 Subjects

Participants were from the Alzheimer’s Disease Neuroimaging Initiative archives. ADNI (adni.loni.usc.edu) was launched in 2003 as a public-private partnership, led by Principal Investigator Michael W. Weiner, MD. The goal of ADNI is to test whether serial magnetic resonance imaging, positron emission tomography, other biological markers, and clinical and neuropsychological assessment can be combined to measure the progression of mild cognitive impairment and early Alzheimer’s disease. For up-to-date information, see www.adni-info.org. Ethics approval was obtained from each study site and all research participants provided written informed consent. ADNI inclusion criteria are listed in Appendix A.

For the purpose of this study, subjects had to have a clinical assessment that included a neuropsychiatric inventory (NPI)(Cummings, 1997) within 3 months (92 days) of their MRI acquisition. Subject age and sex were also obtained from the ADNI database. All subjects were further selected based on quality control of WMH and deformation-based morphometry (DBM) measurements. After quality control of image registrations and WMH segmentation, there were 1131 subjects. Of these subjects, 661 had matching NPI scores (Figure 1). Longitudinal data included between 1-6 follow-up visits over 1-5 years.

**Figure 1.**
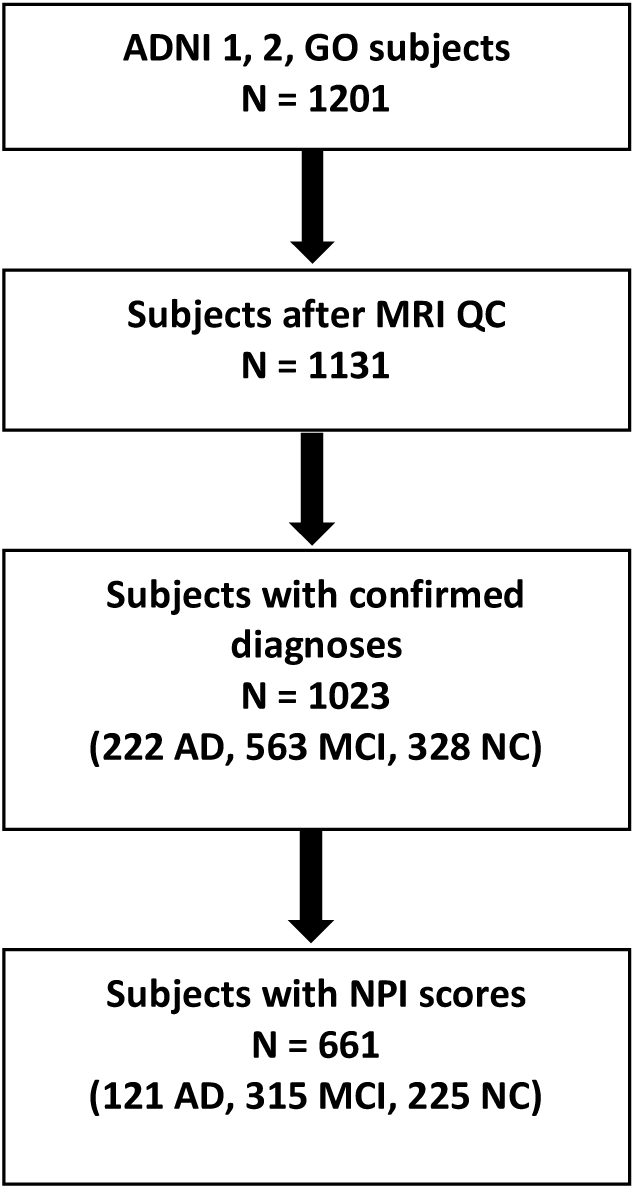
Flowchart of subject inclusion. WMH=white matter hyperintensity, NPI=neuropsychiatric inventory, AD=Alzheimer’s disease, MCI=mild cognitive impairment, NC=normal control.

### 2.2 NPS Assessment

The NPI is primarily used in AD and other dementias to determine behavioural changes that may have occurred since the onset of illness. The following 12 neuropsychiatric domains are assessed by a caregiver/informant: delusions, hallucinations, agitation/aggression, depression/dysphoria, anxiety, elation/euphoria, apathy/indifference, disinhibition, irritability/lability, motor disturbance, night-time behaviour, and appetite/eating. Total symptom scores (0-12) were calculated by multiplying symptom frequency (1-4) and severity (1-3) scores.

### 2.3 Factor Analysis

Based on the factor analysis model from Aalten et al.(Aalten, Verhey, & Boziki, 2007), scores for four neuropsychiatric subsyndromes were calculated: hyperactivity, psychosis, affective symptoms, and apathy. The following NPI symptoms were included in each subsyndrome: hyperactivity (agitation, euphoria, disinhibition, irritability, aberrant motor behaviour), psychosis (delusions, hallucinations, sleep), affective symptoms (depression, anxiety) and apathy (apathy, appetite).

### 2.4 Vascular Risk Factors

Subjects considered to have vascular risk factors were those who reported having cardiovascular history including myocardial infarction, angina, history of smoking, hypertension, stroke, high cholesterol or those on vascular medication. Presence of one or more vascular risk factors was scored as “1” while absence of any vascular risk factors was scored as “0”. We assessed the effect of mild vascular risks as subjects with severe cerebrovascular risk factors were excluded by ADNI.

### 2.5 MRI Processing

T1-weighted, T2-weighted (T2w) and Fluid Attenuated Inversion Recovery (FLAIR) MRI scans were used in this study. All MRI scans were pre-processed with these steps: i) denoising(Manjon, Coupe, Marti-Bonmati, Collins, & Robles, 2010), ii) intensity inhomogeneity correction(Sled, Zijdenbos, & Evans, 1998), and iii) intensity normalization(Fonov et al., 2011). T2w/PDw, and FLAIR scans were rigidly co-registered to the T1w scans(D. Louis Collins, Neelin, Peters, & Evans, 1994). T1w scans were registered to the ADNI template in stereotaxic space(D L Collins & Evans, 1997). Concatenating the two transformations, the other contrasts were also registered to the ADNI template. Using a previously validated automatic WMH segmentation technique and a library of manual segmentations from ADNI dataset, WMHs were segmented for all longitudinal timepoints(Dadar et al., 2018; Dadar, Maranzano, et al., 2017; Dadar, Pascoal, et al., 2017). Quality of segmentations was assessed and verified by an expert (MD). Total WMH volumes (cm3) were calculated, normalized for head size and log-transformed to achieve normal distribution.

DBM analysis was performed using MNI MINC tools. Pre-processed images were linearly (using a 9-parameter rigid registration)(D. Louis Collins et al., 1994) and then non-linearly warped(D. Louis Collins, Holmes, Peters, & Evans, 1995) to the ADNI template. The local deformation obtained from the non-linear transformations was used as a measure of tissue expansion or atrophy. Mean DBM values were calculated for 116 GM ROIs based on the Automated Anatomical Labeling (AAL) atlas(Tzourio-Mazoyer et al., 2002). Results were corrected for multiple comparisons using False Discovery Rate (FDR), thresholded at .05.

### 2.6 Statistical Analysis

Longitudinal mixed-effects models were used to assess the association of total WMH volume and regional GM atrophy with changes in NPI symptoms. The four NPI factors (hyperactivity, psychosis, affective symptoms, apathy) were used as dependent variables. Three different sets of models were run for each of the 116 AAL atlas structures to assess the relationship between NPI, GM, and WMH.

Model 1: NPI Factor ∼ 1+ Age + Sex + Cohort + GM + (1|ID) + (1|Modality)

Model 2: NPI Factor ∼ 1+ Age + Sex + Cohort + WMH + (1|ID) + (1|Modality)

Model 3: NPI Factor ∼ 1+ Age + Sex + Cohort + GM + WMH + (1|ID) + (1|Modality)

Models 1 and 2 were run to assess the relationship between NPI factors with GM atrophy and WMH volume independently. Model 3 includes both GM and WMH as predictors, to assess which one is a more significant contributor to NPS. We adopt this strategy instead of a more complex mediation analysis due to the relatively small number of subject time points available in comparison to the relatively large number of anatomical structures to be tested.

Age, WMH load (denoted in models as WMH), and mean GM DBM values in different ROIs (denoted in models as GM) were used as continuous predictors. Sex (male versus female) and Cohort (normal control versus MCI or AD) were used as categorical predictors. Subjects (denoted by ID) and the modality of the sequences used for segmenting the WMH (FLAIR versus T2w) were used as categorical random variables in all models. All continuous variables were z-scored prior to analysis. All results were corrected for multiple comparisons using FDR, thresholded at 0.05. Models were fitted using fitlme in MATLAB version R2017b.

To examine the relationship between GM and WMH volumes, linear correlations were run between regional GM DBM values and total WMH volumes. One-way ANOVA with Tukey post-hoc tests were used to measure group differences in age, MMSE and NPI total scores.

## 3. Results

Study participants included AD (N=121), MCI (N=315) and normal controls (N=225).

Subjects with MCI were significantly younger than normal controls and AD participants (p<.01). Ages ranged from 55-94 years. There were no differences in sex between groups. NPI total scores were significantly different between groups, with higher values for MCI and AD groups, respectively (p<.001).

Combining all groups, age was not significantly associated with NPI total scores (*β*=.02, p=.50) (Figure 2, left) but was significantly associated with greater WMH load (*β*_*NC*_=0.43, *β*_*MCI*_=0.51, *β*_*AD*_=0.61, p<.001) (Figure 2, center). Relationships with age were examined using the following models: a) NPI Factor ∼ 1+ Age + Cohort + (1|ID) + (1|Modality) and b) WL ∼ 1+ Age + Cohort + (1|ID) + (1|Modality).

**Figure 2.**
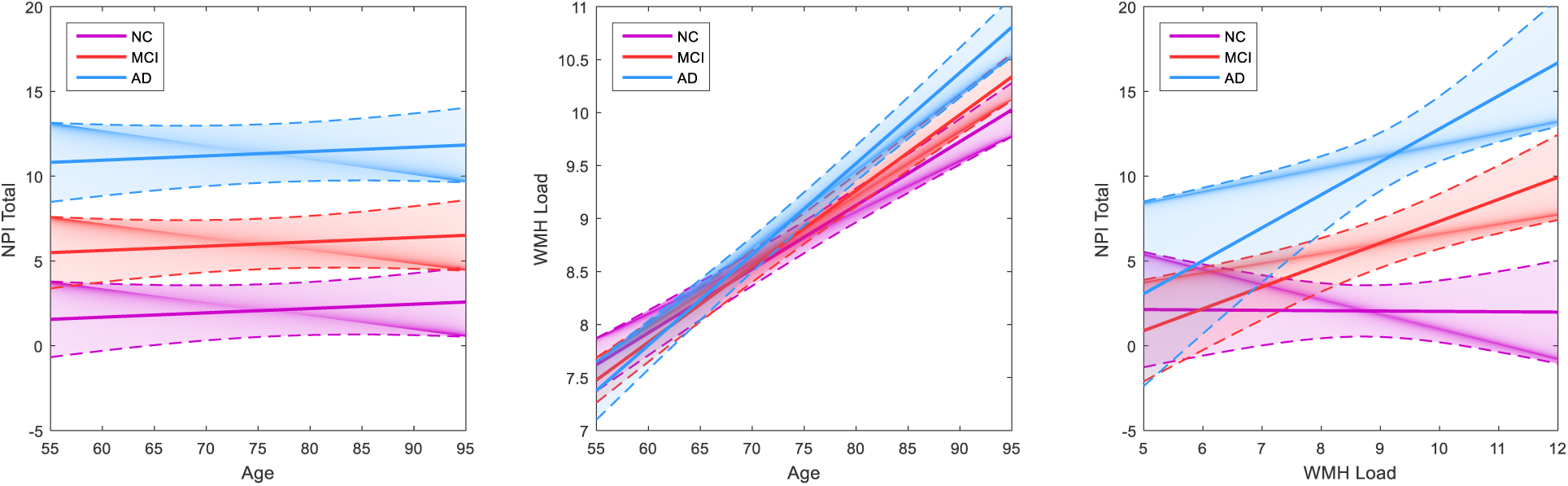
Graphs showing (A) age not associated with NPI total scores, (B) age associated with white matter hyperintensity load, and (C) the relationship between NPI total scores and WMH load across AD, MCI and normal control cohorts.

WMH load was a significant contributor to NPI total scores in MCI and AD cohorts, but not in the normal controls (*β*_*NC*_=0.00, p=.95; *β*_*MCI*_=0.16, p=.02; *β*_*AD*_=0.24, p=.008) (Figure 2, right). This relationship was observed using the following model: NPI Factor ∼ 1+ WL * Cohort + (1|ID) + (1|Modality). WMH load was significantly related to lower DBM GM values after FDR correction (regions listed in Appendix B, Table I).

NPS were divided into hyperactivity, psychosis, affective and apathy subsyndromes for further analysis. DBM analysis identified ROIs where GM was significantly associated with NPS subsyndromes, p<.05 after FDR correction (Table 2, Figure 3). The relationship between NPS with GM atrophy in these ROIs and WMH volume were examined independently. Longitudinal mixed-effects models found AD and MCI cohorts and GM of specific regions to be significantly associated with all four NPS subsyndromes, while sex and age were not significantly associated with any NPS factors (Model 1). WMH volume was also significantly associated with all four NPS subsyndromes (Model 2, Figure 4). Both GM and WMH were then included together as predictors, to assess which one is a more significant contributor to NPS. WMH volume was found to be a significant contributor to NPS subsyndromes while GM DBM values did not remain significant after correcting for multiple comparisons (Model 3). Please see Appendix B, Table II for results of models 1 and 3 before FDR correction, and Table III for results for model 2.

**Table 1.** Subject Demographics.

| Variable | Controls<br>(n=225) | MCI<br>(n=315) | AD<br>(n=121) | *P* |
| --- | --- | --- | --- | --- |
| Age (mean, SD) | 75.58 (6.65) | 73.21 (7.93) | 75.77 (7.38) | 0.0002*<sup>†</sup> |
| Sex (M/F) | 127/98 | 203/112 | 82/39 | 0.147 |
| MMSE at baseline<br>(mean, SD) | 29.14 ± 1.11 | 27.95 ± 1.86 | 21.39 ± 4.62 | p<.0001*<sup>†‡</sup> |
| NPI Total Score at<br>baseline (mean, SD) | 1.17 ± 2.45 | 5.34 ± 7.99 | 8.51 ± 9.32 | p<.001*<sup>†‡</sup> |

MCI=Mild cognitive impairment; AD=Alzheimer's disease
\*p<.01 between AD and MCI
<sup>†</sup>p<.01 between Controls and MCI
<sup>‡</sup>p<.01 between Controls and AD

**Table 2.** List of ROIs where GM DBM values were significantly associated with NPS subsyndromes from results of longitudinal mixed-effects models, before FDR correction. ROIs are from the AAL atlas.

|  | Hyperactivity | Psychosis | Affective | Apathy |
| --- | --- | --- | --- | --- |
| ROIs | L middle frontal gyrus | L Heschl's gyrus | L Cerebellum Crus2 | L Frontal middle gyrus |
|  |  | L Cerebellum 7b |  | L Frontal superior gyrus |
|  | L Cerebellum Crus2 | L Cerebellum Crus2 |  | L Cerebellum Crus2 |
|  | R Inferior Temporal gyrus | R Middle Temporal gyrus |  | R Frontal superior gyrus |
|  | R Cerebellum 7b | R Cerebellum Crus2 |  | R Temporal middle gyrus |
|  | R Cerebellum Crus2 |  |  | R Precentral gyrus |

L=left hemisphere, R=right hemisphere, ROI=region of interest

**Figure 3.**
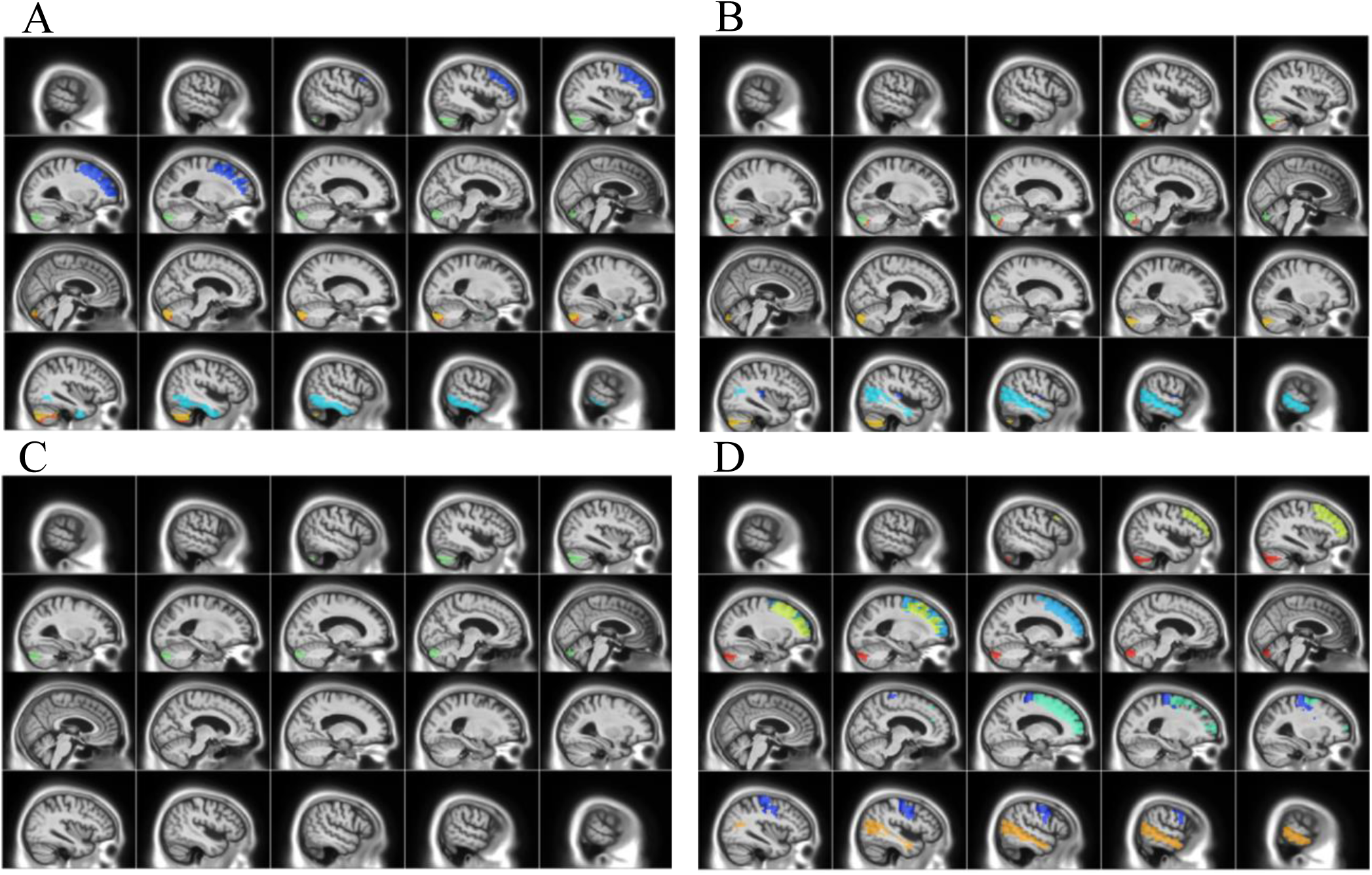
Brain regions where GM DBM values were significantly associated with NPS subsyndrome scores.

**Figure 4.**
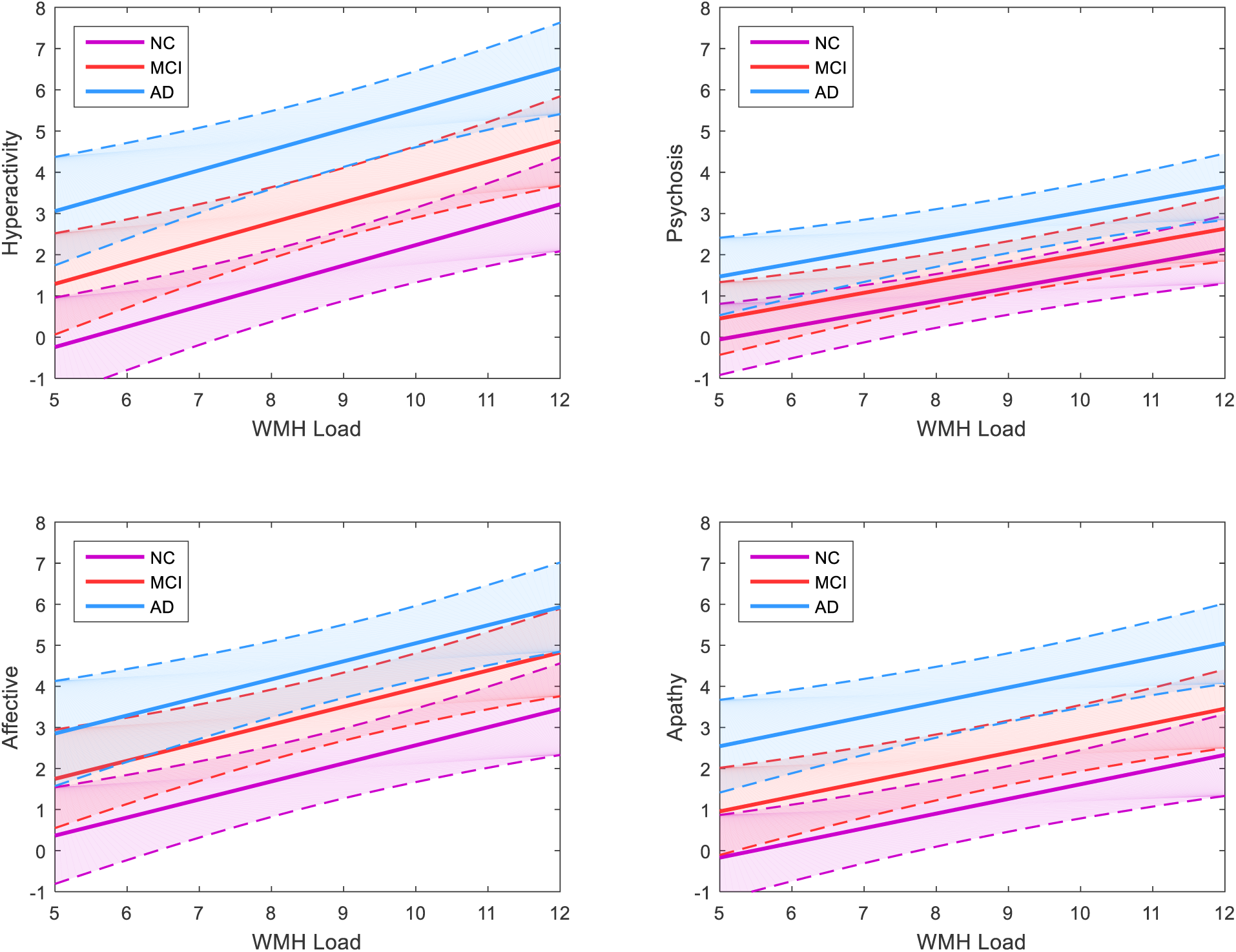
Graphs showing longitudinal mixed-effects model results for the relationship between WMH load volume and hyperactivity, psychosis, affective and apathy NPS subsyndrome scores in AD, MCI and normal controls.

There was a marginal correlation between total vascular risk factors and WMH load (r=0.077, p=.048). Inclusion of vascular risk factors into the model had a significant but minimal contribution to NPS psychosis and affective subsyndrome scores (please see Appendix B, Table IV).

Age is not significantly associated with NPI total scores in any cohort (left). All groups show a positive significant association of age with higher WMH load (middle), and AD and MCI cohorts show a positive significant association between higher WMH load and greater NPI total scores (right). WMH=white matter hyperintensity, NPI=neuropsychiatric inventory, AD=Alzheimer’s disease, MCI=mild cognitive impairment, NC=normal control.

GM ROIs significantly associated with A) hyperactivity B) psychosis C) affective and D) apathy subsyndrome scores. Colors indicate AAL atlas brain regions associated with subsyndromes (please see Appendix B, Table I for complete list of ROIs).

Modelling of mean WMH load (log transformed) on NPS subsyndrome scores shows relationship between higher NPS scores associated with increased WMH load. WMH=white matter hyperintensity, AD=Alzheimer’s disease, MCI=mild cognitive impairment, NC=normal control.

## 4. Discussion

Our findings indicate a significant association of GM atrophy and WMH burden to worsening NPS over time in MCI and AD, with WMH having the greater contribution to NPS. Higher number of vascular risk factors had a significant although minimal effect on worsening psychosis and affective NPS subsyndromes.

The AD cohort showed a higher incidence of NPS compared to MCI and normal control cohorts. This finding is consistent with a majority of studies that have found NPS to be more frequent in AD compared to MCI populations(Geda et al., 2008). Moreover, NPI scores were associated with WMH load in AD and MCI but not with age (despite an age range of 54-94 years in our subjects), suggesting a relationship with disease severity. Although atrophy, as in our study, is associated with NPS and partially explains the increased NPI scores in AD, our findings suggest that WMH burden may also explain increased NPS scores as we found a significant association between WMH load and all four NPS subsyndromes. Our results that WMH burden increases with age are consistent with previous findings(Gouw et al., 2011). Compared to controls, WMH load was greater in AD and MCI subjects and although WMH load is known to increase during healthy aging, this effect appeared to be exaggerated in our AD group as their age was similar to our control group.

Cerebrovascular disease (CVD) is a major cause of WMH. In the current study, there was only a small correlation between total vascular risk factors and WMH load. This is likely because we did not examine the severity of these risk factors and were only able to examine mild cerebrovascular effects as people with significant cerebrovascular risk factors were excluded from ADNI1(Petersen, 2010). There may also be other contributors to the WMH including non-ischemic causes such as enlarged Virchow-Robin spaces(Gouw et al., 2011).

Frontotemporal GM atrophy was implicated in NPS. Psychosis was associated with lower GM DBM values in the right middle temporal gyrus and left Heschl’s gyrus. Heschl’s gyrus, which contains the primary auditory cortex, is involved in acoustic processing while the middle temporal region has been implicated in a number of functions, including semantic memory, audition and language(Xu et al., 2015). Functional connectivity studies have shown that a fronto-temporal language network is impaired in individuals with psychosis(Solé-Padullés et al., 2017), while others have also implicated frontoparietal networks(Baker et al., 2014). Anor et al.(Anor et al., 2017) reported greater delusions with right frontal WMH volume in AD. Delusions have been associated with later Braak staging V/VI(Ehrenberg et al., 2018) suggesting that these symptoms are more likely to present in the late stages of AD. GM atrophy and WMH burden in these regions may contribute to the disruption of connectivity in frontotemporal networks.

The right inferior temporal region was also related to hyperactivity and has been implicated in attention deficit hyperactivity disorder (ADHD)(Zhao et al., 2017). Apathy and hyperactivity were related to lower GM DBM values in the left middle frontal cortex and apathy alone was associated with bilateral superior frontal regions. Apathy is frequently associated with abnormalities in the prefrontal cortex and the basal ganglia in volumetric and functional studies(Bruen et al., 2008; Hahn et al., 2013). Ballarini et al.(Ballarini et al., 2016) identified a relationship between hyperactivity and NPI scores with increased glucose metabolism in predominantly left frontal and limbic structures while apathy scores were negatively correlated with bilateral orbitofrontal and dorsolateral frontal cortex metabolism. Bilateral involvement of brain regions with apathy, and involvement of left hemisphere regions with hyperactivity, was similarly observed in this study. The authors also noted bilateral frontal and limbic involvement with affective symptoms while we did not find any significant regions associated with this subsyndrome.

There were no forebrain or midbrain regions associated with the affective subsyndrome in this study. However, previous reviews of functional and structural MR studies have identified regions associated with depression including the dorsal and medial prefrontal cortex, dorsal and ventral anterior cingulate cortex, orbitofrontal cortex and insula(Pandya, Altinay, Malone, & Anand, 2012), and anxiety with frontoparietal networks and the left ventrolateral prefrontal and superior temporal regions(Picó-Pérez, Radua, Steward, Menchón, & Soriano-Mas, 2017). Our lack of findings may be due to inhomogeneity in the affective subsyndrome group or greater complexity in the regions and networks involved in depression and anxiety. Another possibility is that the NPI is an informant-based questionnaire and so we do not know the actual patient state.

Atrophy in AD follows a pattern from temporal and limbic regions, to frontal and eventually occipital areas of the brain(Thompson et al., 2003). Here, we observed psychosis, apathy and hyperactivity NPS associated with lower GM DBM values in the temporal and frontal regions in subjects with MCI or early AD, suggesting deficits may result from atrophy associated with an early neurodegenerative process.

We found reduced GM DBM values of the left cerebellum crus2 associated with all NPS subsyndromes and the right crus2 with psychosis and hyperactivity. Left cerebellar region 7b was associated with psychosis while the right cerebellar region 7b was associated with hyperactivity. These regions, part of the inferior semilunar lobule, were related to emotion in a study on pain processing(Diano et al., 2016). The authors identified three clusters involved in processing pain: cluster V (vermis IV-V, hemispheres IV-V-VI) was associated with sensory-motor areas, cluster VI (hemisphere VI, crus1, crus2) with cognition and cluster VII (7b, crus1, crus2) with emotion. The role of these regions in NPS may further depend on their functional connectivity with other brain regions and networks involved in regulating emotion and behaviour.

We found WMH burden to be a greater contributor to NPS compared to GM atrophy. The effect of WMH on NPS may be independent of other factors, however it is more likely that a combination of underlying AD and vascular neuropathology contributes to NPS. Neurofibrillary tangles (NFTs) are common in subcortical structures early in AD pathogenesis and these regions (i.e. brainstem, hypothalamic nuclei) are known to be involved in the regulation of NPS(Ehrenberg et al., 2018). The presence of subcortical WMH suggests that SVD may further contribute to damage caused by accumulation of NFTs. SVD may even be involved early in the disease process by accelerating amyloid deposition in AD as a result of impaired perivascular drainage of amyloid-β(Grimmer et al., 2012). Similarly, a recent study found that higher volume of ante-mortem WMH volume predicted greater odds of having autopsy-confirmed AD neuropathology, suggesting SVD may contribute to the severity and progression of AD(Alosco et al., 2018). They identified brain regions where lower GM DBM values were related to higher WMH load, indicating atrophy and CVD pathology may act synergistically in contributing to NPS in MCI and AD.

The importance of WMH in MCI and early AD suggests WMH load is not secondary to AD pathology but an important contributor to NPS during early stages of disease. Neuropsychiatric function may be based on connectivity between brain regions, and therefore the disruption of connectivity by ischemic lesions could have a greater effect than GM atrophy. In addition, genetic and environmental factors may contribute to NPS development and expression.

This study was limited by ADNI protocol exclusion of subjects with psychotic features, agitation or behavioural problems within 3 months prior to screening as this would interfere with study compliance. It is also possible that participants with higher NPS may have been lost to follow-up. In addition, a Hachinski Ischemic Score (HIS) cutoff of 4 was used as part of ADNI selection criteria. As HIS scores greater than 7 suggest vascular involvement, this cutoff may have excluded patients with severe WMH burden making the results here even more compelling. By combining symptoms into subsyndromes, we may have masked relationships with specific NPS such as depression and irritability. Also, although there is evidence for clustering of NPS, there is large variation and overlap of symptoms. Moreover, this study did not take into consideration the use of medication to treat NPS, such as antidepressants and stimulants, and these could have affected the frequency and severity of symptoms reported in the NPI. Although using a large longitudinal dataset improved the robustness of our findings, longer follow-up times would provide a better understanding of the trajectory of NPS in relation to changes in WMH over time. The presence of NPS at the MCI stage suggests the potential benefit of early identification of these symptoms as indicators of disease progression or to monitor response to treatment.

This study identified WMH load as a significant contributor to NPS in AD and MCI using longitudinal data from the large multi-center database of the Alzheimer’s Disease Neuroimaging Initiative. CVD is a common comorbidity of AD and is thought to be the major underlying cause of WMH. Since CVD has a number of modifiable risk factors, such as lowering blood pressure and controlling diabetes and hypercholesterolemia, interventions that aim to reduce vascular disease may prove beneficial in preventing NPS.

## 5. Acknowledgements

We thank all ADNI participants for the generous contribution of their time.

Data collection and sharing for this project was funded by the Alzheimer’s Disease Neuroimaging Initiative (ADNI) (National Institutes of Health Grant U01 AG024904) and DOD ADNI (Department of Defense award number W81XWH-12-2-0012). ADNI is funded by the National Institute on Aging, the National Institute of Biomedical Imaging and Bioengineering, and through generous contributions from the following: AbbVie, Alzheimer’s Association; Alzheimer’s Drug Discovery Foundation; Araclon Biotech; BioClinica, Inc.; Biogen; Bristol-Myers Squibb Company; CereSpir, Inc.; Cogstate; Eisai Inc.; Elan Pharmaceuticals, Inc.; Eli Lilly and Company; EuroImmun; F. Hoffmann-La Roche Ltd and its affiliated company Genentech, Inc.; Fujirebio; GE Healthcare; IXICO Ltd.; Janssen Alzheimer Immunotherapy Research & Development, LLC.; Johnson & Johnson Pharmaceutical Research & Development LLC.; Lumosity; Lundbeck; Merck & Co., Inc.; Meso Scale Diagnostics, LLC.; NeuroRx Research; Neurotrack Technologies; Novartis Pharmaceuticals Corporation; Pfizer Inc.; Piramal Imaging; Servier; Takeda Pharmaceutical Company; and Transition Therapeutics. The Canadian Institutes of Health Research is providing funds to support ADNI clinical sites in Canada. Private sector contributions are facilitated by the Foundation for the National Institutes of Health (www.fnih.org). The grantee organization is the Northern California Institute for Research and Education, and the study is coordinated by the Alzheimer’s Therapeutic Research Institute at the University of Southern California. ADNI data are disseminated by the Laboratory for Neuro Imaging at the University of Southern California.

## Appendix A

The following ADNI inclusion criteria were used for AD: memory complaint verified by a study partner, abnormal memory function measured by delayed recall on the Wechsler Memory Scale Logical Memory II, Mini-Mental State Examination (MMSE) scores between 20-26 (inclusive) and a Clinical Dementia Rating (CDR) scale score of 0.5 or 1.0. Subjects had mild AD and met NINCDS/ADRDA criteria for probable AD1. Criteria for MCI was based on report of memory concern through self-report or report by an informant or clinician, and abnormal memory function measured by delayed recall on the Wechsler Memory Scale Logical Memory II, MMSE score of 24-30 (inclusive) and CDR score of 0.5 (Memory box score at least 0.5). MCI inclusion was dependent on having preserved activities of daily living and no signs of dementia.

## Appendix B: Supplementary Tables

**Table I.** AAL atlas regions where lower GM DBM values are significantly related to higher WMH load, after FDR correction (p<.05). Regions indicated in bold are unique to the left or right hemisphere.

| Left Hemisphere Regions |  | Right Hemisphere Regions |  |
| --- | --- | --- | --- |
| Precentral gyrus | Inferior Occipital | Precentral gyrus | Postcentral gyrus |
| Superior Frontal | Fusiform | Superior Frontal | Superior Parietal |
| Superior Orbital Frontal | Postcentral gyrus | Superior Orbital Frontal | Inferior Parietal |
| Middle Frontal | Superior Parietal | Middle Frontal | Supramarginal |
| Middle Orbital Frontal | Inferior Parietal | Middle Orbital Frontal | Angular |
| Inferior Frontal (pars opercularis) | Supramarginal | Inferior Frontal (pars opercularis) | Precuneus |
| **Inferior Frontal (pars triangularis)** | Angular | Supplementary Motor Area | Paracentral lobule |
| Inferior Frontal (pars orbitalis) | Precuneus | **Olfactory** | Putamen |
| Supplementary Motor Area | Paracentral lobule | **Superior Medial Frontal** | Pallidum |
| **Rectus** | Putamen | Middle Cingulum | Thalamus |
| **Insula** | Pallidum | Posterior Cingulum | Heschl's gyrus |
| **Anterior Cingulum** | Thalamus | Parahippocampal gyrus | **Superior temporal** |
| Middle Cingulum | Heschl's gyrus | Amygdala | **Superior Temporal Pole** |
| Posterior Cingulum | Middle Temporal | Calcarine | Middle Temporal |
| Parahippocampal gyrus | Middle temporal pole | Cuneus | Middle temporal pole |
| Amygdala | Inferior temporal | Lingual gyrus | Inferior temporal |
| Calcarine | Cerebellum Crus 1 | Superior Occipital | Cerebellum Crus 1 |
| Cuneus | Cerebellum Crus 2 | Middle Occipital | Cerebellum Crus 2 |
| Lingual gyrus | Cerebellum 3,4,5,6,7b,8,9,10 | Inferior Occipital | Cerebellum 3,4,5,6,7b,8,9,10 |
| Superior Occipital | **Cerebellar vermis 3-9** | Fusiform |  |
| Middle Occipital |  |  |  |

**Table II.**
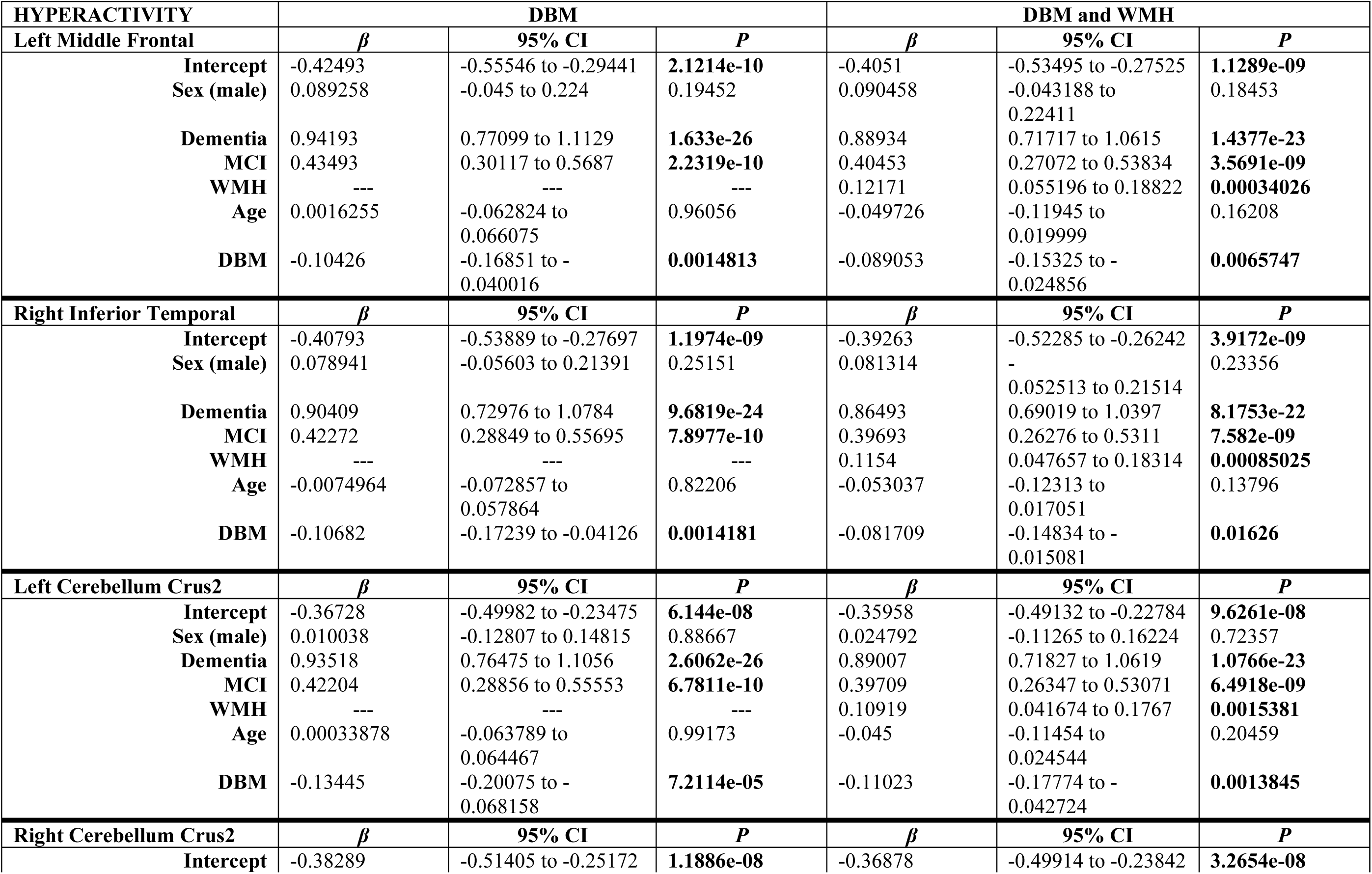
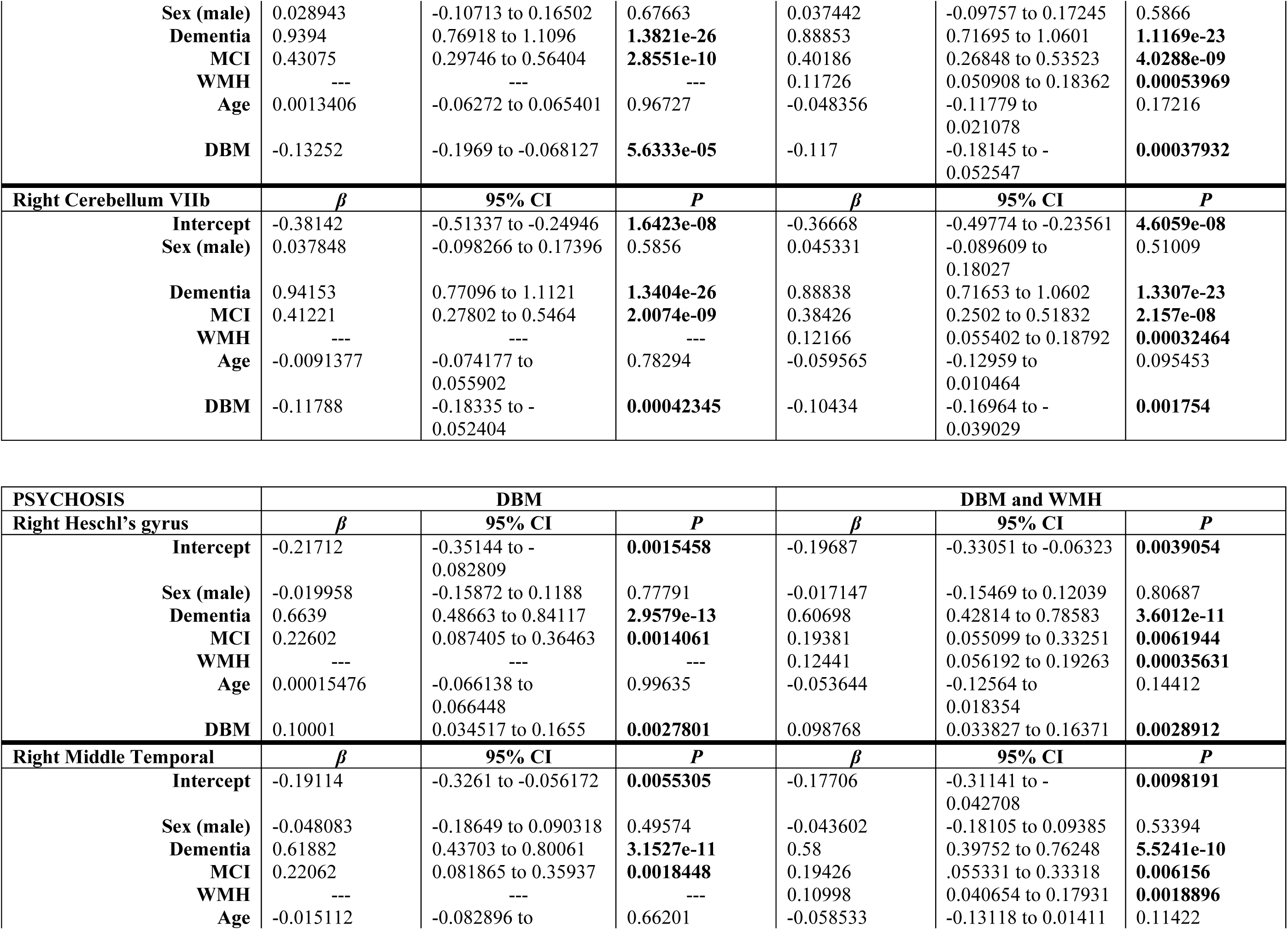
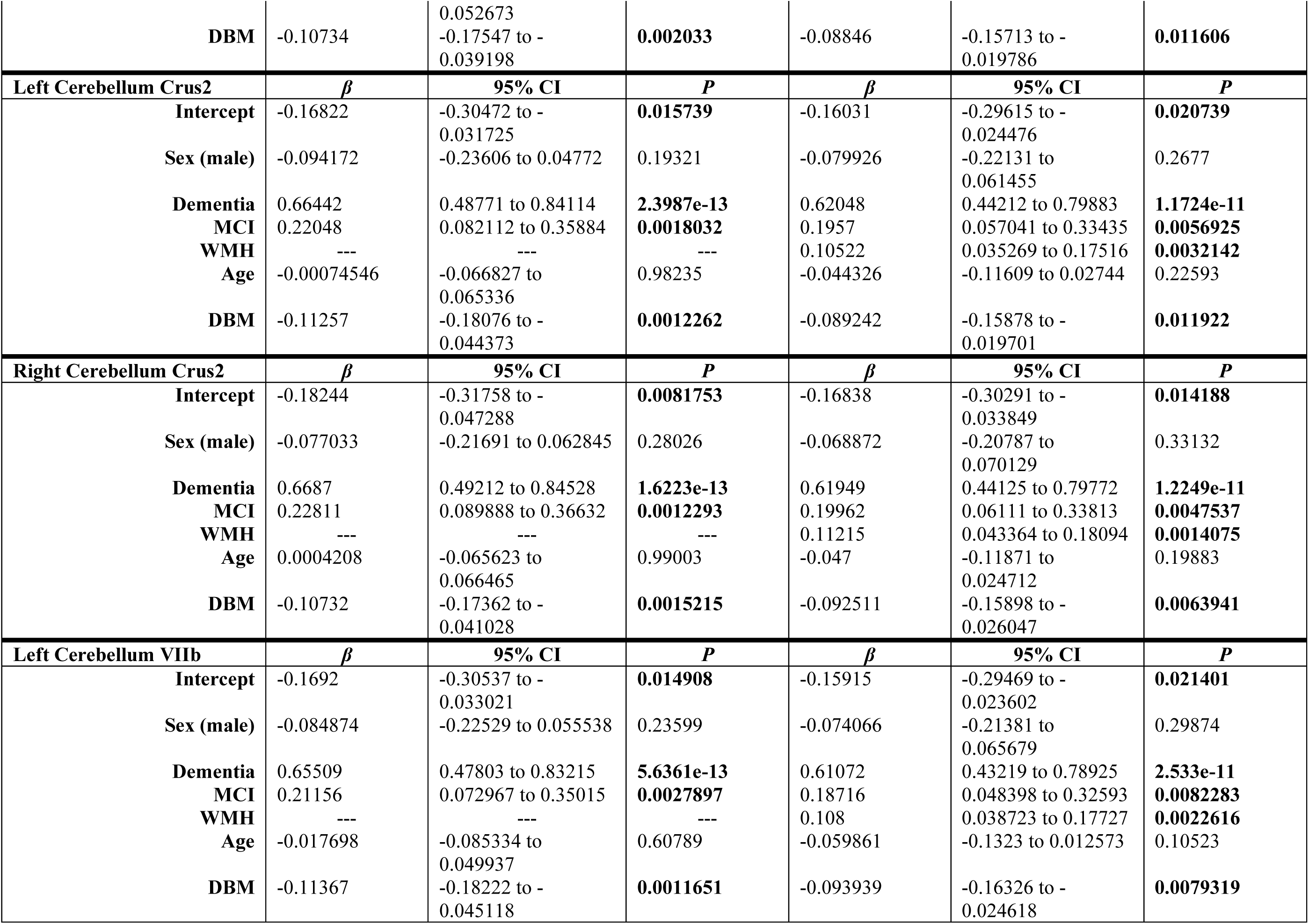
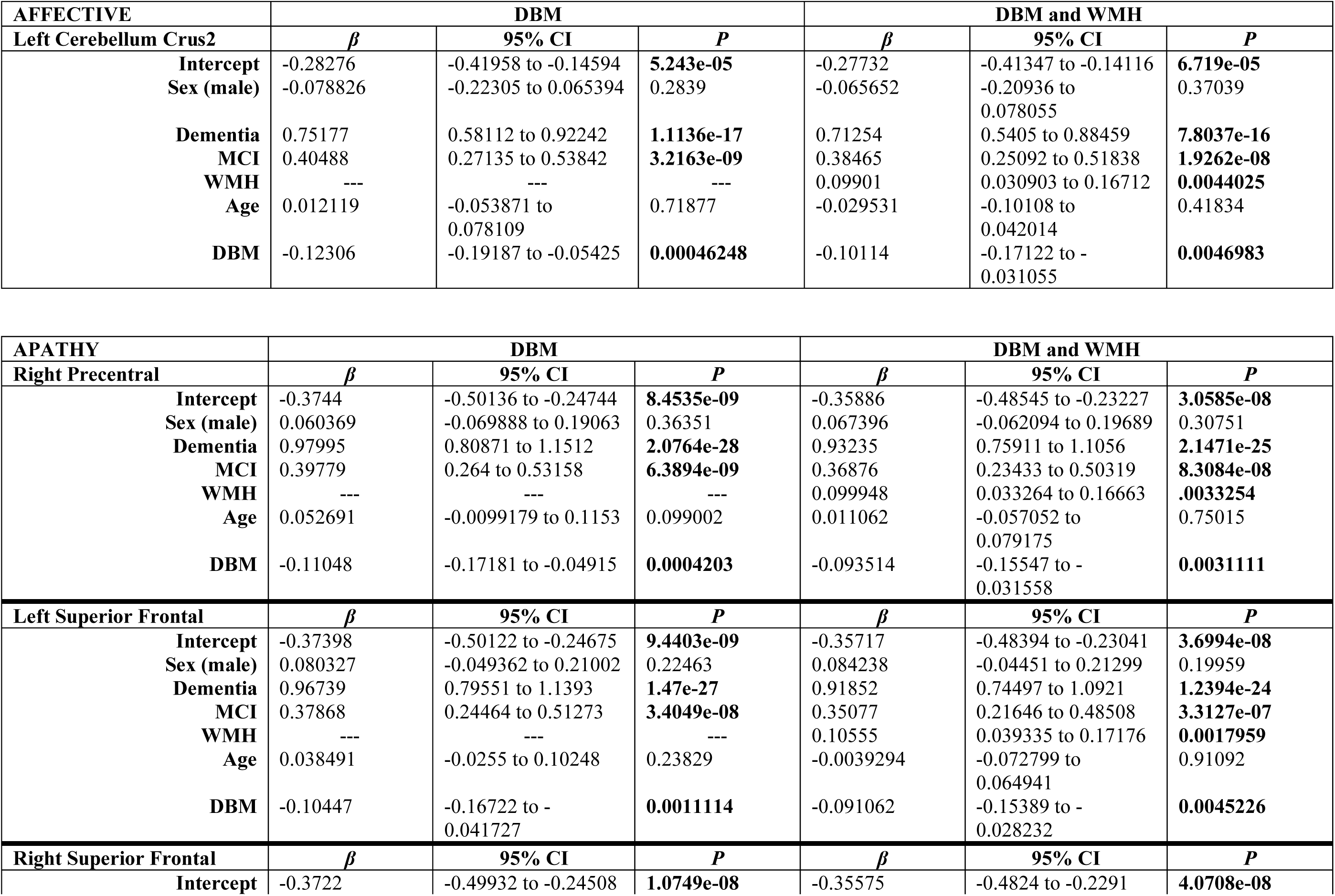
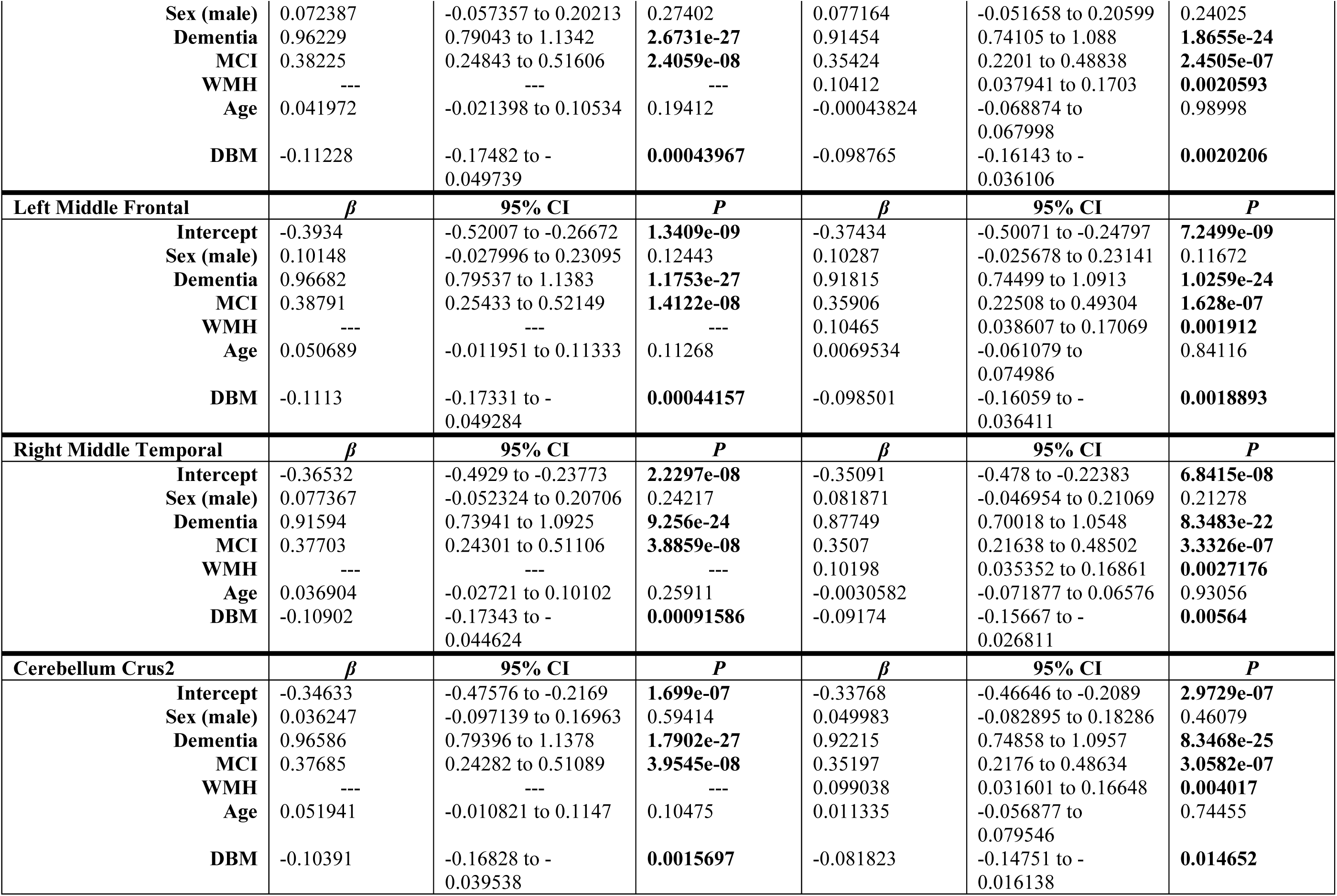
Longitudinal mixed-effects models examining factors contributing to NPS subsyndromes in identified regions of interest. Results with and without WMH included as a factor. Regression coefficients, confidence intervals and p-values are shown for each predictor and NPS subsyndrome. Statistically significant values are indicated in bold, before FDR correction.

**Table III.** Summary of the longitudinal mixed-effects models examining factors contributing to NPS subsyndromes including WMH load as a predictor in the model. Regression coefficients, confidence intervals and p-values are shown for each predictor and NPS subsyndrome. Statistically significant values are indicated in bold.

| Variable | Hyperactivity |  |  | Psychosis |  |  | Affective |  |  | Apathy |  |  |
| --- | --- | --- | --- | --- | --- | --- | --- | --- | --- | --- | --- | --- |
| | $\beta$ | 95% CI | *p* | $\beta$ | 95% CI | *p* | $\beta$ | 95% CI | *p* | $\beta$ | 95% CI | *p* |
| **Intercept** | -0.34 | -0.60 to -0.10 | **0.006** | -0.12 | -0.40 to 0.15 | 0.45 | -0.25 | -0.52 to 0.0003 | 0.05 | -0.31 | -0.59 to -0.02 | **0.03** |
| **WMH load** | 0.12 | 0.06 to 0.18 | **<0.001** | 0.11 | 0.05 to 0.18 | **<0.001** | 0.11 | 0.05 to 0.16 | **<0.001** | 0.12 | 0.06 to 0.18 | **<0.001** |
| **Age** | -0.08 | -0.15 to -0.01 | **0.02** | -0.09 | -0.16 to -0.02 | **0.006** | -0.07 | -0.14 to -0.006 | **0.07** | -0.04 | -0.11 to 0.03 | 0.28 |
| **AD** | 0.92 | 0.75 to 1.09 | **<0.001** | 0.65 | 0.48 to 0.83 | **<0.001** | 0.73 | 0.57 to 0.90 | **<0.001** | 0.93 | 0.76 to 1.10 | **<0.001** |
| **MCI** | 0.42 | 0.29 to 0.56 | **<0.001** | 0.22 | 0.09 to 0.36 | **0.001** | 0.41 | 0.28 to 0.54 | **<0.001** | 0.38 | 0.25 to 0.51 | **<0.001** |
| **Sex (male)** | 0.08 | -0.06 to 0.21 | 0.26 | -0.05 | -0.18 to 0.09 | 0.49 | -0.02 | -0.16 to 0.12 | 0.78 | 0.10 | -0.03 to 0.22 | 0.14 |

NPS=neuropsychiatric symptom; WMH=white matter hyperintensity; MCI=mild cognitive impairment; AD=Alzheimer's disease

**Table IV.**
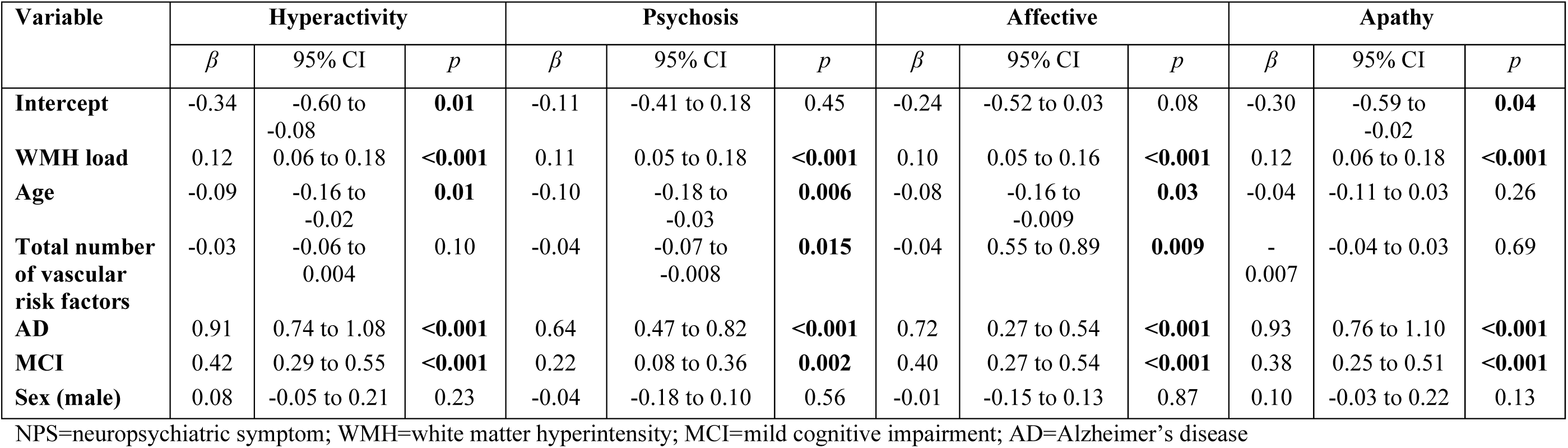
Summary of the longitudinal mixed-effects models examining factors contributing to NPS subsyndromes, with vascular risk factors included as a predictor in the model. Regression coefficients, confidence intervals and p-values are shown for each predictor and NPS subsyndrome. Statistically significant values are indicated in bold.

## Footnotes

1 McKhann G, Drachman D, Folstein M, Katzman R, Price D, Stadlan EM. Clinical diagnosis of Alzheimer’s disease: Report of the NINCDS-ADRDA Work Group* under the auspices of Department of Health and Human Services Task Force on Alzheimer’s Disease. Neurology 1984. doi:10.1212/WNL.34.7.939.

